# The maternal inflammatory proteome during pregnancy and its role in predicting the risk of spontaneous preterm birth

**DOI:** 10.1101/2025.05.19.654577

**Authors:** Oluwanifewa Laleye, Raphaella Jackson, Jiadai Mi, Andrew H Shennan, Natasha Hezelgrave, Alessandra Vigilante, Joan Camunas-Soler, Morten Rasmussen, Rachel M Tribe

## Abstract

**Background:** Spontaneous preterm birth (sPTB) is a significant adverse outcome of pregnancy. Being able to identify and improve the management of those who may be at risk requires robust screening methods. The use of circulating molecular markers provides a promising and non-invasive solution to this problem to allow necessary and successful intervention. The role of inflammation has been continuously demonstrated to play a key role in the onset of sPTB with intrauterine inflammation being a key driver. Here we sought out to explore the inflammatory proteome using a nested case-control approach using samples from pregnant participants in the INSIGHT cohort.

**Objectives:** To explore the maternal blood proteome in the second trimester using the Olink Explore panel to identify inflammatory proteins associated with sPTB and assess their predictive value, both independently and in combination with cell-free RNA (cfRNA).

**Study Design:** We conducted a nested case-control study to investigate inflammatory protein profiles during the second trimester of pregnancy. A total of 138 maternal blood plasma samples were analysed using a targeted proteomic assay quantifying 384 inflammation-related proteins. Differential expression analysis and a LASSO-logistic regression model with Leave-One-Out Cross-Validation (LOOCV) were applied to evaluate the association between inflammatory biomarkers and spontaneous preterm birth (sPTB) outcomes.

**Results:** Using predictive modelling of the maternal blood proteome, 16 inflammation-related proteins were identified as key discriminators of sPTB risk, with proteins such as PGF, COL9A1, CST7, CXCL6 and GALNT3 emerging as major contributors for predicting sPTB risk. Using inflammation-related maternal proteins alone to predict sPTB (<35 weeks) achieved an area under the receiver operating characteristic curve (AUC-ROC) of 0.76 (95% CI: 0.66–0.84). The incorporation of both cfRNA and proteomic data into an integrated model, improved the area under the curve to 0.85 (95% CI: 0.78–0.92). The integrated model highlighted inflammatory biomarkers that are not only implicated in preterm birth but also in essential physiological mechanisms such as placental function, tissue remodelling, and extracellular matrix composition, which are critical to maintaining pregnancy and preventing premature labour.

**Conclusions:** These findings demonstrate that an integrated approach using both cfRNA and proteomic signatures of the second trimester maternal blood plasma yields a more comprehensive biomarker profile for predicting preterm birth risk. This multimodal strategy not only enhances the predictive accuracy but also captures a broader array of biological signals across multiple organ systems. Compared to relying solely on inflammatory proteome markers, this multiomic method offers a deeper molecular characterisation of preterm birth risk in the maternal blood plasma.

## Introduction

Preterm birth, defined as birth occurring before 37 weeks’ gestation, affects an estimated 13.4 million newborns globally each year and remains a significant global health concern (1). For those born prematurely there is an increased risk of poor healthcare outcomes with higher rates of mortality in comparison to infants born at term (1–4). Preterm birth of spontaneous onset (sPTB) is a complex syndrome with many aetiologies (5–8). Inflammation, however, appears to underlie many of the mechanistic routes to sPTB (ref). (9–13). Exploitation of the maternal inflammatory proteome could demonstrate value in the prediction of sPTB risk.

Currently, the identification and management of pregnancies at risk of sPTB is challenged by the lack of a reliable prediction biomarkers to identify pregnancies that may be at risk. The strongest clinical indicator of sPTB risk is a history of PTB, where there is a ∼30% chance of a subsequent preterm birth (14–16). Clinical tools to assess sPTB risk have proved useful particularly for asymptomatic, high-risk women, largely relying on measuring cervical length via ultrasound from 20 weeks’ gestation and/or quantitative fetal fibronectin (qfFN) and (17,18). The QUiPP app, designed for second-trimester use, has successfully utilized obstetric history, cervical length, qfFN, and where relevant labor symptoms to aid decision making in asymptomatic and symptomatic women (19).Recently, however, the commonly used commercial test for qfFN has been discontinued. This has left clinical history and cervical length measurement as the key predictors available for use, with the QuIPP algorithm able to utilise these in the absence of qfFN. Clearly, there is an urgent need for new predictive markers. Identifying early biomarkers in the first or early second trimester will improve early sPTB risk assessment for earlier intervention.

To improve our ability to predict sPTB, we must first deepen our understanding of the underlying biological mechanisms. By studying the proteome, we have the potential to gain valuable insights that can help refine and enhance multiomic models for better risk assessment. Proteomic surveillance for biomarker discovery in sPTB research, has leveraged high throughput methods and technologies, targeting various maternal biofluids such as blood plasma, amniotic fluid, and cervicovaginal secretions (20–27). Predictive modelling methodologies applied to such molecular data show promise in identifying biomarkers that correlate with sPTB risk within these distinct environments. Notably, our group’s previous research has identified 25 cell-free RNA (cfRNA) transcripts with significant predictive potential for sPTB risk (28). The immune system is highly adapted during pregnancy and has been consistently shown that Inflammation is a major of role in the pathophysiology of PTB (9,29,30). Our work aims to discover new biomarkers for the prediction of sPTB utilising inflammation biomarkers. Measurement of inflammatory biomarkers offers a window into how biological changes may contribute to PTB to help identify pregnancies at risk during the earlier stages of pregnancy.

Olink proteomics utilises proximity extension assay (PEA) based technology to perform targeted sequencing for protein targets of interest. Using this technology, we aimed to explore the maternal plasma proteome during the second trimester of pregnancy to identify inflammation-specific markers associated with sPTB. This research aims to expand our understanding of the biology of sPTB and to develop predictive models utilising inflammation-related proteins for identifying sPTB (<35 weeks) risk. This work also aims to build on previous work by integrating these findings with previously identified cfRNA signatures.

## Materials and Methods

### Study Design and Cohort

The study population and sample collection procedures are consistent with those described in the Camunas-Soler et al. 2022 study (28). Specifically, the INSIGHT (Investigation into Biomarkers for the Prediction of Spontaneous Preterm Birth) study, which is a longitudinal observational cohort designed to examine biomarkers in both high-risk and low-risk populations for predicting sPTB (31). Ethical approval for the study was granted by the London City and East Research Ethics Committee (13/LO/1393), and informed consent was obtained from all participants. Women of the INSIGHT cohort were recruited from four tertiary antenatal clinics in the United Kingdom. A nested case-control study design was used and consisted of sPTB cases each being matched with two term birth controls (1 high-risk and 1 low-risk term control). The criteria for matching were based on ethnicity, BMI, smoking status and maternal age. Blood plasma samples were obtained between 16-24 weeks gestation from women having a singleton pregnancy. The categorisation of risk was based on criteria including having one or more of the following: previous sPTB or late miscarriage (16–37 weeks’ gestation), a history of cervical surgery, the presence of uterine anomaly, or a cervical length of <25 mm as determined by transvaginal ultrasound. If no risk-factors for sPTB were identified at the time of enrolment, these women were characterised as low-risk. Women with multiple pregnancies, significant fetal anomalies, membrane rupture, or vaginal bleeding at recruitment were excluded from both risk groups. sPTB classification was assigned if they experienced spontaneous labour or preterm premature rupture of membranes leading to delivery before 37 weeks. To strengthen our study design and enhance discovery power, we intentionally excluded individuals who delivered between 35 and 37 weeks, to allow a clearer distinction between term and preterm births. No preeclampsia cases were included in case or control groupings. Full details on participant recruitment and selection criteria are available in (28).

### Sample Collection

Blood samples were collected between 16–24 weeks gestation based on due dates estimated by first-trimester ultrasound. Samples were collected in EDTA tubes and processed within four hours of collection. After centrifugation (2500 g, 10 min, 4°C), plasma was aliquoted and stored at −80°C until analysis. A total of 176 blood plasma samples assayed for inflammatory proteins. From this initial group, a subset of 138 samples with matching cfRNA data were selected to develop and validate a predictive model for sPTB. This subset comprised 46 samples from sPTB cases and 92 samples from term birth controls, encompassing 46 high-risk and 46 low-risk pregnancies.

### Inflammation-related Proteomic Assay using the Olink Inflammation 384 panel

Targeted proteomic analysis was performed using the Olink Inflammation panel which enables the identification of 368 inflammatory markers of interest. Olink uses proximity extension assay (PEA) technology that involves each protein being detected by a matched pair of antibodies tethered to DNA oligonucleotides. Antibody binding allows oligonucleotides to be in proximity to target protein, hybridise and extend which is then quantified by real-time qPCR to establish protein amounts. Olink uses an arbitrary NPX (Normalised Protein eXpression) as a unit of measurement where an increase of 1 NPX corresponds to a doubling of the relative protein concentration (log2 scale). Final protein quantification is presented using NPX where a high value corresponds to a high protein concentration. The complete list of proteins (and full names) in the Olink Inflammation Panel is provided in Supplementary Table 1.

### Proteomic Assay Quality Control

Assays for proteins BID, MGLL and BCL211 were removed due to not meeting Olink’s quality control criteria. Proteins that were measurable above the limit of detection in at least 25% of the samples were included (Supplementary Table 2), leaving 368 proteins for further analysis and model development.

### Statistical Analysis

A nonparametric Mann-Whitney U test assessed significance across cohort demographics, such as birth outcome groups and age, as well as pregnancy-related details, including obstetric history. For ethnicity, a Chi-square test evaluated significance. Differential protein expression between sPTB cases and term birth controls was visualised using volcano plots. The ‘Olink® Analyze’ package facilitated the exploration of differentially expressed proteins (DEPs) between preterm and term birth outcomes, which employs a Welch’s two-sample t-test for differential protein expression analysis.

### Predicative Model Development

To identify inflammation-related proteins with potential to predict sPTB risk, we developed two predictive models. The first model incorporated only inflammation-related proteomic markers. Protein selection for this model was guided by differential expression analysis using t-tests, comparing sPTB cases to term birth controls. We performed t-tests under two conditions: first, comparing all sPTB cases to all term controls, and second, comparing all sPTB cases to only low-risk term controls. The inclusion threshold for the model was proteins with a p-value <0.05, resulting in a combined list of 27 proteins. Both models included characteristics regarding maternal age and BMI at booking and gestational age at sample collection.

Biomarker selection and model performance utilised a leave-one-out cross-validation (LOOCV) approach. In each LOOCV iteration, an L1-Lasso logistic regression model was used to facilitate feature selection (32). Receiver operating characteristic (ROC) curve confidence intervals (CI) were determined through bootstrapping. The second model combined the selected 27 proteomic markers with 25 previously identified cfRNA transcripts (28), using the same analysis workflow as the first model.

## Results

### Cohort Demographics

A total of 138 maternal blood plasma samples taken from the second trimester were successful analysed for this study. Of these samples, 46 were from pregnancies that ended in a sPTB (gestation <35 weeks) and 92 were from pregnancies that ended in a term birth (gestation ≥37 weeks). 46 of these term pregnancies were classified as being at high risk of preterm birth at the time of their enrolment into the INSIGHT study. Table 1 provides descriptive information regarding the characteristics of the cohort; there were no missing data. Women who had a previous sPTB event or were classified as high risk at enrolment had significantly higher rates (p<0.001^1^) of sPTB than those who did not. Gestational age at sampling for sPTB cases and term controls were statistical different (p-value 0.003^2^). Other demographic characteristics including BMI, maternal age at enrolment and ethnicity showed no significant differences between sPTB cases and term birth controls groups.

**Figure 1.**
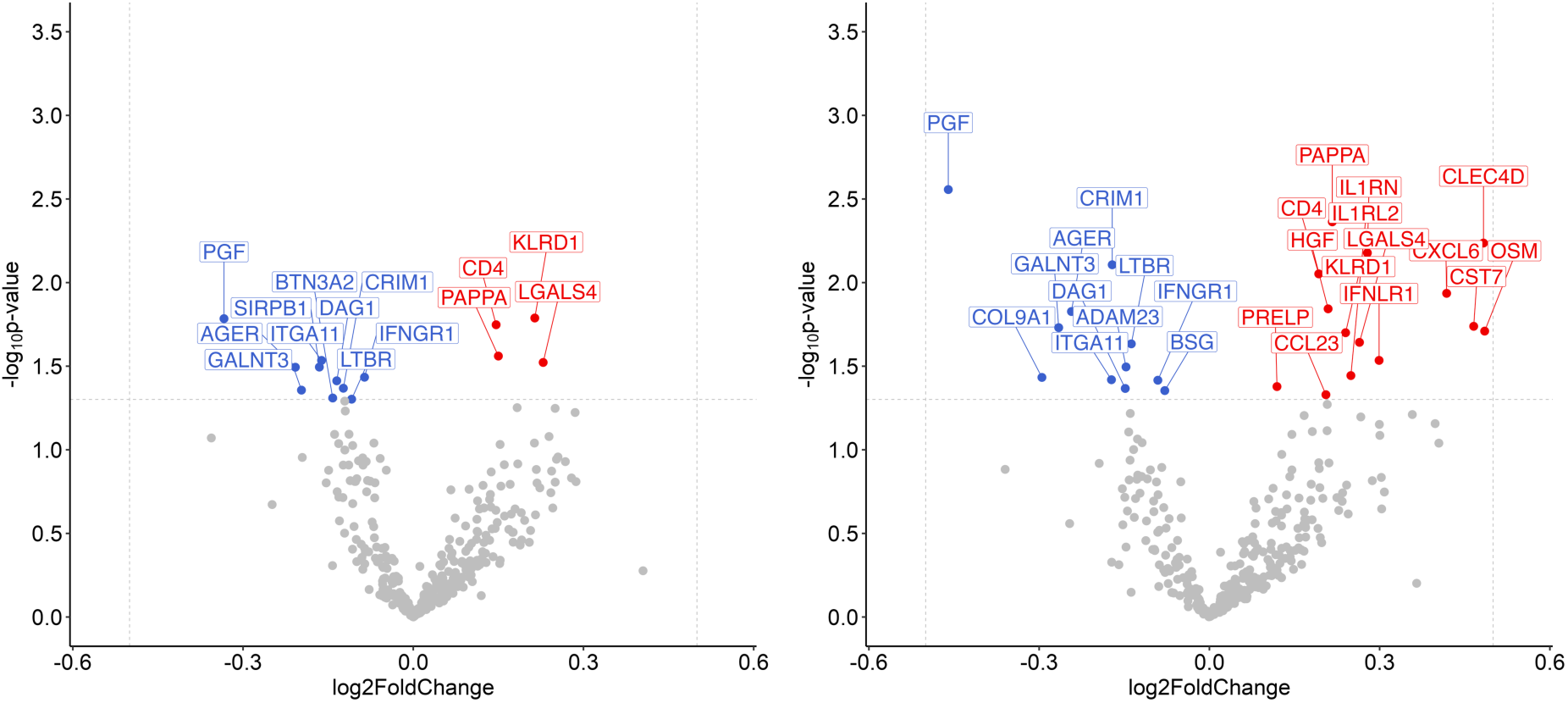
Differential expression of inflammation-related proteins in maternal plasma taken between 16-24 weeks of gestation from participants prior to a spontaneous preterm birth (sPTB, < 35 weeks, cases) and term birth (> 37 weeks, controls). Red indicates upregulated proteins and Blue indicates downregulated proteins. A) Volcano plot analysis of differentially expressed proteins (DEPs) of 368 inflammatory proteins for all sPTB cases (n=46) and all term birth controls (both high-risk and low-risk term controls, n=92). B) Volcano plot analysis of DEPs of 368 inflammatory proteins for all sPTB cases (n=46) and only low-risk term birth controls (n=46).

**Figure 2.**
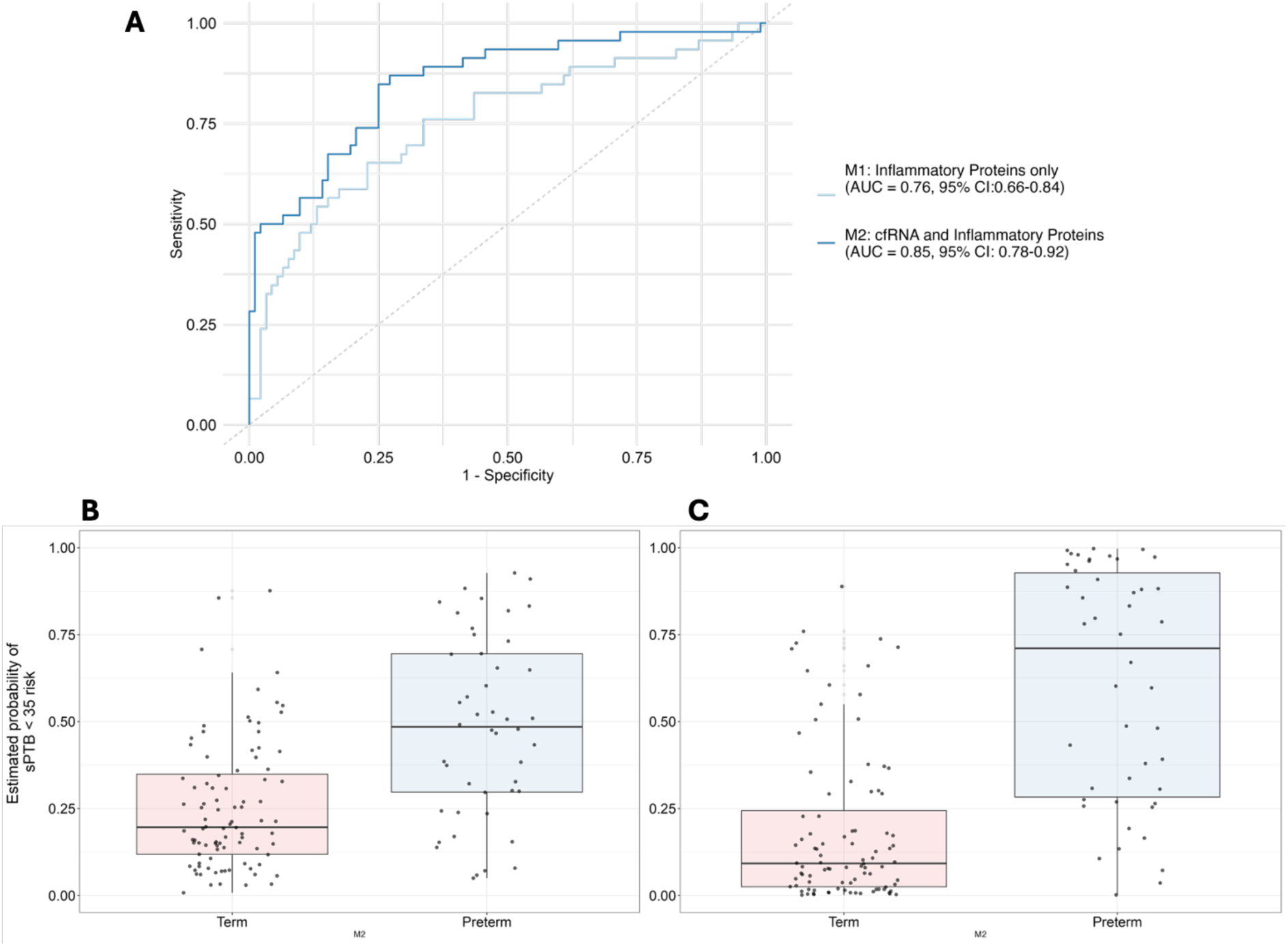
Model performance for the prediction of spontaneous preterm birth (sPTB, <35 weeks’ gestation) risk when using only maternal plasma inflammatory proteomic markers and integrating with cfRNA transcripts. A) ROC performance for sPTB (<35 weeks’ gestation) when the model is using only inflammation-related proteomic markers and the ROC performance for the integrated model using both inflammatory proteins and cfRNA. B) The probability assigned by the inflammatory proteomic model to each maternal plasma sample. A value of 0 corresponds to 0% probability and a value of 1 corresponds to 100% probability of a sample being classified as a sPTB case. C) The probability assigned by the integrated model to each sample. sPTB cases = 46 (target 0), Term birth controls = 92 (target 1). M1 – Model1, M2 – Model 2.

**Table 1.** Descriptive characteristics of study cohort demographics, preterm birth risk factors at enrolment, and length of gestation at delivery. P-value derived from a Mann Whitney U test for all groups except for ethnicity where it was derived using a Chi-Square test.

|  |  | Overall | Preterm | Term | P-Value |
| --- | --- | --- | --- | --- | --- |
| **n** |  | 138 | 46 | 92 |  |
| **Outcome, n (%)** | **Preterm** | 46 (33.3) | 46 (100.0) |  | <0.001 |
|  | **Term** | 92 (66.7) |  | 92 (100.0) |  |
| **Previous sPTB 37 weeks, n (%)** | **No** | 105 (76.1) | 28 (60.9) | 77 (83.7) | 0.006 |
|  | **Yes** | 33 (23.9) | 18 (39.1) | 15 (16.3) |  |
| **Ethnicity, n (%)** | **Asian** | 17 (12.3) | 5 (10.9) | 12 (13.0) | 0.957 |
|  | **Black** | 42 (30.4) | 14 (30.4) | 28 (30.4) |  |
|  | **Other** | 10 (7.2) | 4 (8.7) | 6 (6.5) |  |
|  | **White** | 69 (50.0) | 23 (50.0) | 46 (50.0) |  |
| **Low risk at enrolment, n (%)** | **No** | 87 (63.0) | 41 (89.1) | 46 (50.0) | <0.001 |
|  | **Yes** | 51 (37.0) | 5 (10.9) | 46 (50.0) |  |
| **Gestation at delivery (weeks), mean (SD)** |  | 35.3 (6.4) | 27.7 (5.6) | 39.1 (1.2) | <0.001 |
| **BMI (kg/m<sup>2</sup>), mean (SD)** |  | 26.4 (5.8) | 26.9 (6.0) | 26.1 (5.8) | 0.462 |
| **Maternal age (years), mean (SD)** |  | 33.1 (5.8) | 33.5 (6.0) | 33.0 (5.7) | 0.610 |
| **Primigravida, n (%)** | **No** | 111 (80.4) | 43 (93.5) | 68 (73.9) | 0.012 |
|  | **Yes** | 27 (19.6) | 3 (6.5) | 24 (26.1) |  |
| **Gestational age at sample collection (weeks), mean (SD)** |  | 19.1 (1.9) | 18.3 (1.8) | 19.5 (1.8) | 0.003 |

### Differences in the Inflammatory Proteome by sPTB Outcome and Risk Type

Two analyses of differential expression were performed to compare profiles of inflammatory protein expression for case and control groups to identify potential early biomarkers of sPTB. The first analysis compared samples from pregnancies that went on to have a sPTB *versus* samples from all women who went on to have term births. The second analysis compared samples from women who went on to have a sPTB *versus* samples from women who went on to have term births who were also classified as being at low risk of preterm birth at enrolment.

Comparing sPTB samples to all term samples, there were 14 differentially expressed proteins (p<0.05^3^), of these four were upregulated and 10 were downregulated in sPTB samples compared to term samples (Figure 1A). Comparing sPTB samples to only low-risk term samples we found there were 25 differentially expressed proteins (p<0.05), of these 14 were upregulated and 11 were downregulated (Figure 1B). Across these two comparisons a total of 27 different inflammatory proteins were identified as being differentially expressed with eight proteins being common across both analyses.

**Figure 3.**
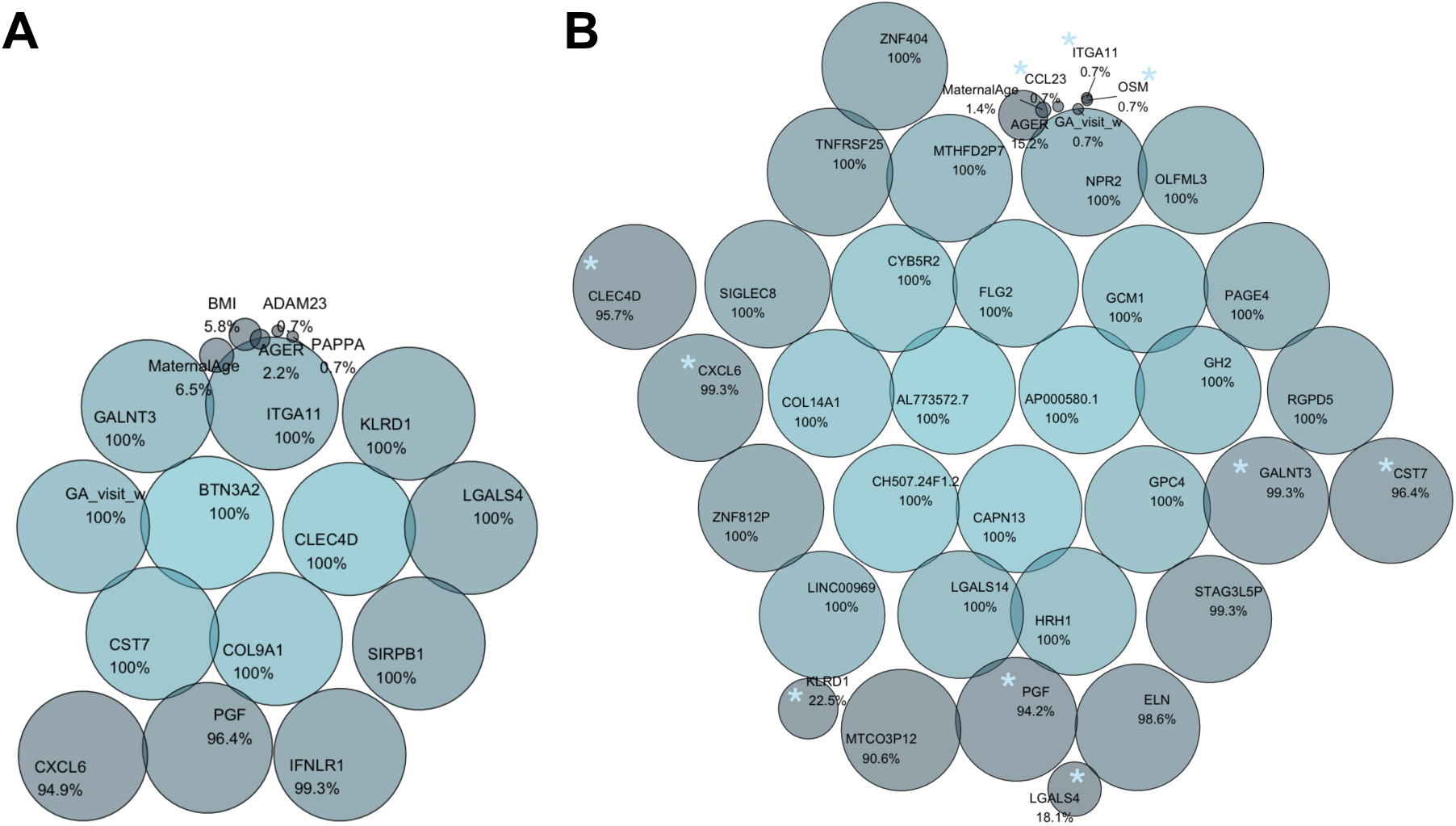
Bubbleplot visualisation of model-derived feature importance for (a) the proteomic model and (b) the combined proteomic and cell-free RNA model (proteomic markers are highlighted by blue asterisk (*) symbol). Each circle represents one molecular marker, with both the size and colour intensity reflecting its proportional contribution (%) to the final predicted outcome during feature discovery. Marker names and proportional frequencies are labelled within each bubble.

### Predictive Modelling of sPTB Risk

To identify candidate inflammation-related proteins predictive of early sPTB risk (<35 weeks), an initial logistic regression model was developed and validated using a LOOCV framework. From an initial set of 27 proteins of interest, 16 proteins emerged as potential predictors of sPTB risk. Gestation at sample collection contributed greatly to the model, with maternal age and BMI having smaller contributions (Figure 3a). The proteomic-only model achieved an AUC-ROC of 0.76 (95% CI: 0.66–0.84) (Figure 2a). Figure 3 highlights the frequency of protein usage in the model, with KLRD1, PGF, COL9A1, SIRPB1, AGER, LGALS4, CST7, IFNLR1, GALNT3, BTN3A2 and ITGA11 being particularly predictive of sPTB risk. The integrated cfRNA-proteomic model, which combined 25 cfRNA transcripts with the 27 proteins of interest, achieved a considerably higher AUC-ROC of 0.85 (95% CI: 0.78–0.92) (Figure 2a). All cfRNA transcripts consistently contributed to the predictive capability of the model. Among inflammation-related proteins, CXCL6, GALNT3, PGF, CST7, CLEC4D, KLRD1, LGALS4 and AGER were meaningful predictors in this multiomic model. For this model, maternal characteristics had little influence (Figure 3b). The probability of sPTB risk (<35 weeks’ gestation), was computed with the distribution of probabilities shown in Figure 2B and Figure 2C for the respective models. The second model, which used an integrated omics approach, demonstrated a greater separation in sPTB risk probabilities compared to the proteomic-only model.

## Discussion

This study explored the potential of the maternal plasma inflammatory proteome alone, and in combination with previously identified cfRNAs (28) to predict sPTB risk. Along with identifying inflammatory proteins associated with sPTB, the key finding was that an integrated ‘omics approach improved the accuracy of predicting sPTB risk <35 weeks of gestation compared to using proteomic markers in isolation. This new model offers a more robust tool for early sPTB risk detection.

Recent efforts in predicting sPTB risk have utilised proteomic markers across various biological fluids, including maternal blood, amniotic fluid, and cervicovaginal fluid, but very few have been translated into widespread routine clinical use (33–37) (20,22,23).

Multiomic approaches, integrating proteomics with other ‘omic data like cfRNA, addresses the growing realisation that combining multi-biological pathways enhances predictive power and improves early identification of at-risk pregnancies (24,26). Studies incorporating machine learning models across multiomic datasets show promising results (24,26), highlighting the need for a multifaceted approach that accounts for the complexity of biological systems.

Notably, a recent longitudinal study suggested that plasma proteomic models outperform transcriptomic approaches in predicting sPTB, when sampling between 27-33 weeks of gestation. This highlighted the potential of using even earlier pregnancy quantification of the proteome for sPTB prediction (26),(10). In this earlier window, proteomic models were not as good as cfRNA models alone (AUC 0.80, (28)).

Our integrated model, which analysed plasma markers between 16-24 weeks’ gestation, achieved an AUC-ROC of 0.85. The model also identified several proteins of interest early in gestation. We considered the role of maternal BMI, gestation at sample collection, and maternal age, as potential confounding factors. However, neither BMI nor maternal age had a significant impact on the utility of both models for predicting sPTB risk.

Both models highlighted some unique proteins which may have mechanistic relevance. The protein GALNT3, for example, plays a key role in phosphate homeostasis, which fluctuates during pregnancy and may reflect changes in maternal metabolism. Women experiencing threatened preterm delivery have been shown to exhibit serum deficiencies in total calcium, phosphorus, and magnesium, potentially contributing to premature uterine contractions (38,39). The model also incorporated PGF, which is essential for placental development and has established potential as a biomarker for conditions such as preterm preeclampsia (40–42). Several proteins, including GALNT3, PGF, CST7, CLEC4D, KLRD1, OSM, CCL23, AGER and LGALS4, overlapped between the proteomic and multiomic models. The pro-inflammatory protein CXCL6 was found to contribute to both the models developed. CXCL6 is known to be expressed at the maternal-fetal interface and implicated in pregnancy processes (43). CXCL6 is known to be found in the amnionic fluid with higher levels associated with sPTB (44) with elevated levels notable in cases with intra-amniotic infection. However, even in the absence of infection CXCL6 levels remain higher in PTB cases in comparison to those who deliver at term (44) – suggesting a potential role in PTB pathology. Although CXCL6 has not been directly linked to sPTB, its involvement in the immune response during pregnancy warrants further investigation. Additionally, proteins like CLEC4D, LGALS4 AND KLRD1, which play important roles in immune regulation, have not yet been directly associated with sPTB, but further research could elucidate their role in the inflammatory processes that may influence preterm birth risk.

The use of Olink proteomics for quantifying blood-based inflammatory biomarkers holds significant promise for clinical applications. Integrating this high-throughput proteomic technology with cfRNA profiles could potentially serve as a non-invasive and cost-effective predictive tool for identifying women at risk for sPTB. For example, clinical management algorithms such as those used by QUiPP app, which can combine both clinical and quantitative biomarkers (19), could be enhanced by integrating additional markers, such as those discussed here. These additional biomarkers may significantly improve sPTB risk prediction.

This study expands the scope of preterm birth research by incorporating a multiomic framework, combining the inflammatory proteome with cfRNA transcripts to identify biomarkers of sPTB risk. Understanding how these proteins contribute to health and pathogenesis during pregnancy is essential for identifying complications that may arise and may be associated with inflammatory states. This could also inform more targeted immunotherapy approaches. Future research should continue to explore the integration of different ‘omic layers, particularly focusing on how inflammation interacts with other physiological processes that influence preterm birth.

A strength of this study is the diversity of participants used from the INSIGHT study, which includes both high and low-risk pregnancies. This diversity enhances the generalisability of the findings and ensures that the model can be applied to a wide range of pregnancy phenotypes. Additionally, the analysis of samples taken between 16-24 weeks’ gestation provides valuable insight into the second trimester prior to the development of pregnancy complications. The exploratory nature of the study also allows for a more comprehensive understanding of the maternal inflammatory proteome and its role in preterm birth.

While the Olink Explore 384 Inflammation panels allowed for the assessment of a large number of proteins, which is useful for initial discovery, it was a targeted approach. This means that other potentially important inflammatory markers equally important during pregnancy are not covered in the panel. Additionally, while the inflammatory proteome is likely to be a fundamental factor in preterm birth, it is only one aspect of the complex pathophysiology leading to sPTB. As this study excluded late preterm births (35-37 weeks), future research could validate findings in cohorts that include these cases, providing further insights into the inflammatory profile during this critical period. Future work should also explore additional biological markers and consider broader, untargeted proteomic approaches that may capture a wider range of relevant proteins.

## Conclusion

In conclusion, the integration of cfRNA and protein markers offers a promising non-invasive, multiomic approach for improving the prediction of sPTB risk. While inflammatory proteins alone demonstrate moderate predictive performance, combining them with cfRNA markers improves the model’s accuracy and offers a more reliable tool for early detection. Further validation is needed, but this integrated approach holds promise as a predictive tool for identifying those at risk for preterm birth. As research in multiomics continues to evolve, such approaches may become instrumental in refining predictive models and improving outcomes for both mothers and newborns. Pregnancy is a highly dynamic process, and its complexity may not be fully captured by focusing solely on proteomic inflammation during the second trimester at single time point. If a single test is capable of predicting sPTB outcomes it will likely need to be one which incorporates a multiomic approach so to incorporate dynamic and mechanistic differences in pregnancy and sPTB.

## Acknowledgments

We extend our sincere appreciation to all participants and centers that contributed samples to this study. A special thank you to the CLRN midwifery team, particularly Judy Filmer, Falak Diab, Vicky Robinson, and Debbie Finucane, for their invaluable assistance with recruitment, sample collection, and clinical data acquisition.

We gratefully acknowledge the support of Tommy’s Charity (1060508), the National Institute for Health Research (NIHR) Biomedical Research Centre at Guy’s and St Thomas’ NHS Foundation Trust, and the Rosetrees Trust (298582) (M303-CD1) in facilitating sample collection for the “Insight” study. Additional support was provided by an NIHR Doctoral Research Fellowship (DRF-2013-06-171) awarded to Natasha L. Hezelgrave, as well as funding from Borne (1167073) and Action Medical Research (GN2666).

Joan Camunas-Soler is supported by the Knut and Alice Wallenberg Foundation (Wallenberg Molecular Medicine Fellow), the Swedish Research Council (grant 2021-05109), and the Erling Persson Foundation (Swedish Foundations’ Starting Grant).

## Author Contributions

Author contributions (CRediT): Conceptualization: RMT, MR, JM; Methodology: JCS, MR, RMT, JM, FL; DATA CURATION,: RMT, JCS, MR, FL; FORMAL ANALYSIS: FL, RJ, JCS; Investigation: JCS, JM, FL, RJ; Visualization: FL, JCS, MR, RMT; Funding Acquisition: RMT, AHS, NH; Project Administration: RMT, JM; Supervision: AV, RMT, RJ; Writing original draft: FL, RMT, RJ; Writing – reviewing and editing: FL, RJ, JM, AHS, NH, AV, RMT,JCS, MR

## Conflicts of Interest

The coauthors affiliated with Mirvie are inventors of patented applications that cover the detection, diagnosis, or treatment of pregnancy complications and/or have an equity interest in Mirvie. All the cohort contributors were compensated for sample collection and/or shipping.

## Footnotes

1 Mann-Whitney U

2 Pearson’s chi-squared test

3 Welch’s t-test

## Notes

### Summary of Updates

This manuscript has been revised to include figures in the correct format for Figure 1, Figure 2 and Figure 3

