## Supplemental Table 1 for "The maternal inflammatory proteome during pregnancy and its role in predicting the risk of spontaneous preterm birth"

Supplementary Table 1. Names of proteins included in the Olink Explore 384 Inflammation Panel

| OlinkID | UniProt | Gene Name | Protein Name |
| --- | --- | --- | --- |
| OID20435 | O43707 | ACTN4 | Alpha-actinin-4 |
| OID20645 | P00813 | ADA | Adenosine deaminase |
| OID20651 | O75077 | ADAM23 | Disintegrin and metalloproteinase domain-containing protein 23 |
| OID20755 | Q9UHX3 | ADGRE2 | Adhesion G protein-coupled receptor E2 |
| OID20756 | Q15109 | AGER | Advanced glycosylation end product-specific receptor |
| OID20786 | O00468 | AGRN | Agrin |
| OID20658 | O00253 | AGRP | Agouti-related protein |
| OID20512 | P30838 | ALDH3A1 | Aldehyde dehydrogenase, dimeric NADP-preferring |
| OID20437 | Q9NP70 | AMBN | Ameloblastin |
| OID20492 | Q9BXJ7 | AMN | Protein amnionless |
| OID20740 | Q15389 | ANGPT1 | Angiopoietin-1 |
| OID20726 | Q9UKU9 | ANGPTL2 | Angiopoietin-related protein 2 |
| OID20703 | Q9BY76 | ANGPTL4 | Angiopoietin-related protein 4 |
| OID20583 | P50995 | ANXA11 | Annexin A11 |
| OID20452 | P19801 | AOC1 | Amiloride-sensitive amine oxidase [copper-containing] |
| OID20554 | Q9NZN5 | ARHGEF12 | Rho guanine nucleotide exchange factor 12 |
| OID20432 | P27540 | ARNT | Aryl hydrocarbon receptor nuclear translocator |
| OID20446 | Q5T4W7 | ARTN | Artemin |
| OID20760 | Q9UII2 | ATP5IF1 | ATPase inhibitor, mitochondrial |
| OID20582 | O15169 | AXIN1 | Axin-1 |
| OID20780 | P15291 | B4GALT1 | Beta-1,4-galactosyltransferase 1 |
| OID20505 | O14867 | BACH1 | Transcription regulator protein BACH1 |
| OID20594 | Q8NDB2 | BANK1 | B-cell scaffold protein with ankyrin repeats |
| OID20513 | O43521-2 | BCL2L11 | BIM (Bcl-2 Interacting Mediator of cell death) |
| OID20558 | P11274 | BCR | Breakpoint cluster region protein |
| OID20517 | P55957 | BID | BH3 Interacting Domain Death Agonist |
| OID20626 | P35613 | BSG | Basigin |
| OID20713 | Q7KYR7 | BTN2A1 | Butyrophilin subfamily 2 member A1 |
| OID20540 | P78410 | BTN3A2 | Butyrophilin subfamily 3 member A2 |
| OID20654 | P02745 | C1QA | Complement C1q subcomponent subunit A |
| OID20555 | P42575 | CASP2 | Caspase-2 |
| OID20668 | P51671 | CCL11 | Eotaxin |
| OID20655 | Q99616 | CCL13 | C-C motif chemokine 13 |
| OID20745 | Q92583 | CCL17 | C-C motif chemokine 17 |
| OID20671 | P78556 | CCL20 | C-C motif chemokine 20 |
| OID20686 | O00585 | CCL21 | C-C motif chemokine 21 |
| OID20765 | O00626 | CCL22 | C-C motif chemokine 22 |
| OID20693 | P55773 | CCL23 | C-C motif chemokine 23 |
| OID20772 | O00175 | CCL24 | C-C motif chemokine 24 |
| OID20674 | O15444 | CCL25 | C-C motif chemokine 25 |
| OID20546 | Q9Y258 | CCL26 | C-C motif chemokine 26 |
| OID20569 | Q9NRJ3 | CCL28 | C-C motif chemokine 28 |
| OID20610 | P10147 | CCL3 | C-C motif chemokine 3 |
| OID20695 | P13236 | CCL4 | C-C motif chemokine 4 |
| OID20523 | P80098 | CCL7 | C-C motif chemokine 7 |
| OID20709 | P29279 | CCN2 | CCN family member 2 |
| OID20647 | O95971 | CD160 | CD160 antigen |
| OID20603 | P41217 | CD200 | OX-2 membrane glycoprotein |
| OID20595 | Q8TD46 | CD200R1 | Cell surface glycoprotein CD200 receptor 1 |
| OID20637 | P20273 | CD22 | B-cell receptor CD22 |
| OID20628 | Q9BZW8 | CD244 | Natural killer cell receptor 2B4 |
| OID20680 | Q5ZPR3 | CD276 | CD276 antigen |
| OID20584 | P01730 | CD4 | T-cell surface glycoprotein CD4 |
| OID20724 | P25942 | CD40 | Tumor necrosis factor receptor superfamily member 5 |
| OID20616 | P29965 | CD40LG | CD40 ligand |
| OID20692 | P09326 | CD48 | CD48 antigen |
| OID20716 | P19256 | CD58 | Lymphocyte function-associated antigen 3 |
| OID20649 | P30203 | CD6 | T-cell differentiation antigen CD6 |
| OID20599 | P32970 | CD70 | CD70 antigen |
| OID20635 | P40259 | CD79B | B-cell antigen receptor complex-associated protein beta chain |
| OID20565 | Q01151 | CD83 | CD83 antigen |
| OID20607 | Q9UIB8 | CD84 | SLAM family member 5 |
| OID20754 | Q4KMG0 | CDON | Cell adhesion molecule-related/down-regulated by oncogenes |
| OID20667 | Q15517 | CDSN | Corneodesmosin |
| OID20548 | Q3KPI0 | CEACAM21 | Carcinoembryonic antigen-related cell adhesion molecule 21 |
| OID20441 | Q9UPV0 | CEP164 | Centrosomal protein of 164 kDa |
| OID20771 | Q9BU40 | CHRDL1 | Chordin-like protein 1 |
| OID20617 | Q07065 | CKAP4 | Cytoskeleton-associated protein 4 |
| OID20721 | P12532 | CKMT1A_CKMT1B | Creatine kinase U-type, mitochondrial |
| OID20573 | Q9UMR7 | CLEC4A | C-type lectin domain family 4 member A |
| OID20547 | Q8WTT0 | CLEC4C | C-type lectin domain family 4 member C |
| OID20609 | Q8WXI8 | CLEC4D | C-type lectin domain family 4 member D |
| OID20590 | Q6UXB4 | CLEC4G | C-type lectin domain family 4 member G |
| OID20636 | Q9BXN2 | CLEC7A | C-type lectin domain family 7 member A |
| OID20559 | Q9UDT6 | CLIP2 | CAP-Gly domain-containing linker protein 2 |
| OID20664 | Q9H4D0 | CLSTN2 | Calsyntenin-2 |
| OID20556 | Q9UHC6 | CNTNAP2 | Contactin-associated protein-like 2 |
| OID20550 | P20849 | COL9A1 | Collagen alpha-1(IX) chain |
| OID20738 | Q5KU26 | COLEC12 | Collectin-12 |
| OID20751 | Q6UXH1 | CRELD2 | Protein disulfide isomerase CRELD2 |
| OID20747 | P24387 | CRHBP | Corticotropin-releasing factor-binding protein |
| OID20701 | Q9NZV1 | CRIM1 | Cysteine-rich motor neuron 1 protein |
| OID20742 | P46109 | CRKL | Crk-like protein |
| OID20699 | O75462 | CRLF1 | Cytokine receptor-like factor 1 |
| OID20719 | P09603 | CSF1 | Macrophage colony-stimulating factor 1 |
| OID20491 | P09919 | CSF3 | Granulocyte colony-stimulating factor |
| OID20704 | O76096 | CST7 | Cystatin-F |
| OID20752 | Q99895 | CTRC | Chymotrypsin-C |
| OID20679 | P53634 | CTSC | Dipeptidyl peptidase 1 |
| OID20660 | P43234 | CTSO | Cathepsin O |
| OID20561 | P78310 | CXADR | Coxsackievirus and adenovirus receptor |
| OID20762 | P09341 | CXCL1 | Growth-regulated alpha protein |
| OID20697 | P02778 | CXCL10 | C-X-C motif chemokine 10 |
| OID20464 | P48061 | CXCL12 | C-X-C motif chemokine 12 |
| OID20458 | O95715 | CXCL14 | C-X-C motif chemokine 14 |
| OID20622 | Q6UXB2 | CXCL17 | C-X-C motif chemokine 17 |
| OID20788 | P19876 | CXCL3 | C-X-C motif chemokine 3 |
| OID20613 | P80162 | CXCL6 | C-X-C motif chemokine 6 |
| OID20631 | P10145 | CXCL8 | C-X-C motif chemokine 8 |
| OID20675 | Q07325 | CXCL9 | C-X-C motif chemokine 9 |
| OID20774 | Q14118 | DAG1 | Dystroglycan 1 |
| OID20524 | Q9UN19 | DAPP1 | Dual adapter for phosphotyrosine and 3-phosphotyrosine and 3-phosphoinosit |
| OID20681 | Q9UJU6 | DBNL | Drebrin-like protein |
| OID20579 | Q16698 | DECR1 | 2,4-dienoyl-CoA reductase [(3E)-enoyl-CoA-producing], mitochondrial |
| OID20620 | O00273 | DFFA | DNA fragmentation factor subunit alpha |
| OID20466 | Q13574 | DGKZ | Diacylglycerol kinase zeta |
| OID20627 | O60884 | DNAJA2 | DnaJ homolog subfamily A member 2 |
| OID20712 | Q8NFT8 | DNER | Delta and Notch-like epidermal growth factor-related receptor |
| OID20744 | O43598 | DNPH1 | 2′-deoxynucleoside 5′-phosphate N-hydrolase 1 |
| OID20527 | Q8N608 | DPP10 | Inactive dipeptidyl peptidase 10 |
| OID20525 | Q9UNE0 | EDAR | Tumor necrosis factor receptor superfamily member EDAR |
| OID20698 | P01133 | EGF | Pro-epidermal growth factor |
| OID20572 | Q9GZT9 | EGLN1 | Egl nine homolog 1 |
| OID20625 | Q04637 | EIF4G1 | Eukaryotic translation initiation factor 4 gamma 1 |
| OID20423 | P63241 | EIF5A | Eukaryotic translation initiation factor 5A-1 |
| OID20497 | Q8N8S7 | ENAH | Protein enabled homolog |
| OID20736 | Q9UJA9 | ENPP5 | Ectonucleotide pyrophosphatase/phosphodiesterase family member 5 |
| OID20689 | Q6UWV6 | ENPP7 | Ectonucleotide pyrophosphatase/phosphodiesterase family member 7 |
| OID20743 | P16422 | EPCAM | Epithelial cell adhesion molecule |
| OID20677 | P21709 | EPHA1 | Ephrin type-A receptor 1 |
| OID20522 | P01588 | EPO | Erythropoietin |
| OID20705 | P21860 | ERBB3 | Receptor tyrosine-protein kinase erbB-3 |
| OID20758 | Q9NQ30 | ESM1 | Endothelial cell-specific molecule 1 |
| OID20691 | P25116 | F2R | Proteinase-activated receptor 1 |
| OID20778 | P07148 | FABP1 | Fatty acid-binding protein, liver |
| OID20501 | Q0Z7S8 | FABP9 | Fatty acid-binding protein 9 |
| OID20665 | P48023 | FASLG | Tumor necrosis factor ligand superfamily member 6 |
| OID20564 | P24071 | FCAR | Immunoglobulin alpha Fc receptor |
| OID20639 | Q96LA5 | FCRL2 | Fc receptor-like protein 2 |
| OID20443 | Q96P31 | FCRL3 | Fc receptor-like protein 3 |
| OID20529 | Q6DN72 | FCRL6 | Fc receptor-like protein 6 |
| OID20715 | O95750 | FGF19 | Fibroblast growth factor 19 |
| OID20503 | P09038 | FGF2 | Fibroblast growth factor 2 |
| OID20490 | P12034 | FGF5 | Fibroblast growth factor 5 |
| OID20722 | Q9Y3D6 | FIS1 | Mitochondrial fission 1 protein |
| OID20618 | P68106 | FKBP1B | Peptidyl-prolyl cis-trans isomerase FKBP1B |
| OID20661 | P49771 | FLT3LG | Fms-related tyrosine kinase 3 ligand |
| OID20515 | Q12778 | FOXO1 | Forkhead box protein O1 |
| OID20770 | P19883 | FST | Follistatin |
| OID20782 | O95633 | FSTL3 | Follistatin-related protein 3 |
| OID20463 | Q96DB9 | FXYD5 | FXYD domain-containing ion transport regulator 5 |
| OID20727 | P22466 | GAL | Galanin peptides |
| OID20471 | Q14435 | GALNT3 | Polypeptide N-acetylgalactosaminyltransferase 3 |
| OID20489 | P32456 | GBP2 | Guanylate-binding protein 2 |
| OID20710 | Q9HC38 | GLOD4 | Glyoxalase domain-containing protein 4 |
| OID20641 | P36959 | GMPR | GMP reductase 1 |
| OID20551 | Q9HD26 | GOPC | Golgi-associated PDZ and coiled-coil motif-containing protein |
| OID20663 | P12544 | GZMA | Granzyme A |
| OID20604 | P10144 | GZMB | Granzyme B |
| OID20648 | P14317 | HCLS1 | Hematopoietic lineage cell-specific protein |
| OID20589 | O94992 | HEXIM1 | Protein HEXIM1 |
| OID20656 | P14210 | HGF | Hepatocyte growth factor |
| OID20520 | P01903 | HLA-DRA | HLA class II histocompatibility antigen, DR alpha chain |
| OID20532 | P13747 | HLA-E | HLA class I histocompatibility antigen, alpha chain E |
| OID20615 | P37235 | HPCAL1 | Hippocalcin-like protein 1 |
| OID20575 | P28845 | HSD11B1 | 11-beta-hydroxysteroid dehydrogenase 1 |
| OID20718 | P0DMV8 | HSPA1A | Heat shock 70 kDa protein 1A |
| OID20484 | Q05084 | ICA1 | Islet cell autoantigen 1 |
| OID20578 | Q14773 | ICAM4 | Intercellular adhesion molecule 4 |
| OID20619 | P22304 | IDS | Iduronate 2-sulfatase |
| OID20495 | P01579 | IFNG | Interferon gamma |
| OID20678 | P15260 | IFNGR1 | Interferon gamma receptor 1 |
| OID20506 | Q8IU57 | IFNLR1 | Interferon lambda receptor 1 |
| OID20544 | Q9Y6K9 | IKBKG | NF-kappa-B essential modulator |
| OID20431 | P22301 | IL10 | Interleukin-10 |
| OID20420 | Q13651 | IL10RA | Interleukin-10 receptor subunit alpha |
| OID20696 | Q08334 | IL10RB | Interleukin-10 receptor subunit beta |
| OID20455 | P20809 | IL11 | Interleukin-11 |
| OID20666 | P29460 | IL12B | Interleukin-12 subunit beta |
| OID20486 | P42701 | IL12RB1 | Interleukin-12 receptor subunit beta-1 |
| OID20430 | P35225 | IL13 | Interleukin-13 |
| OID20562 | P40933 | IL15 | Interleukin-15 |
| OID20498 | Q13261 | IL15RA | Interleukin-15 receptor subunit alpha |
| OID20633 | Q14005 | IL16 | Pro-interleukin-16 |
| OID20469 | Q16552 | IL17A | Interleukin-17A |
| OID20477 | Q9P0M4 | IL17C | Interleukin-17C |
| OID20481 | Q8TAD2 | IL17D | Interleukin-17D |
| OID20456 | Q96PD4 | IL17F | Interleukin-17F |
| OID20585 | Q9NRM6 | IL17RB | Interleukin-17 receptor B |
| OID20694 | Q14116 | IL18 | Interleukin-18 |
| OID20652 | Q13478 | IL18R1 | Interleukin-18 receptor 1 |
| OID20488 | P01583 | IL1A | Interleukin-1 alpha |
| OID20427 | P01584 | IL1B | Interleukin-1 beta |
| OID20757 | P27930 | IL1R2 | Interleukin-1 receptor type 2 |
| OID20487 | Q9HB29 | IL1RL2 | Interleukin-1 receptor-like 2 |
| OID20700 | P18510 | IL1RN | Interleukin-1 receptor antagonist protein |
| OID20419 | P60568 | IL2 | Interleukin-2 |
| OID20453 | Q9NYY1 | IL20 | Interleukin-20 |
| OID20422 | Q9UHF4 | IL20RA | Interleukin-20 receptor subunit alpha |
| OID20449 | Q8N6P7 | IL22RA1 | Interleukin-22 receptor subunit alpha-1 |
| OID20440 | Q13007 | IL24 | Interleukin-24 |
| OID20450 | P14784 | IL2RB | Interleukin-2 receptor subunit beta |
| OID20605 | P24001 | IL32 | Interleukin-32 |
| OID20428 | O95760 | IL33 | Interleukin-33 |
| OID20461 | P26951 | IL3RA | Interleukin-3 receptor subunit alpha |
| OID20426 | P05112 | IL4 | Interleukin-4 |
| OID20576 | P24394 | IL4R | Interleukin-4 receptor subunit alpha |
| OID20472 | P05113 | IL5 | Interleukin-5 |
| OID20601 | Q01344 | IL5RA | Interleukin-5 receptor subunit alpha |
| OID20563 | P05231 | IL6 | Interleukin-6 |
| OID20535 | P13232 | IL7 | Interleukin-7 |
| OID20485 | P51617 | IRAK1 | Interleukin-1 receptor-associated kinase 1 |
| OID20577 | Q9NWZ3 | IRAK4 | Interleukin-1 receptor-associated kinase 4 |
| OID20538 | B1AKI9 | ISM1 | Isthmin-1 |
| OID20581 | Q9UKX5 | ITGA11 | Integrin alpha-11 |
| OID20528 | P23229 | ITGA6 | Integrin alpha-6 |
| OID20493 | P18564 | ITGB6 | Integrin beta-6 |
| OID20504 | O43736 | ITM2A | Integral membrane protein 2A |
| OID20494 | P01591 | JCHAIN | Immunoglobulin J chain |
| OID20424 | P05412 | JUN | Transcription factor Jun |
| OID20629 | Q12918 | KLRB1 | Killer cell lectin-like receptor subfamily B member 1 |
| OID20632 | Q13241 | KLRD1 | Natural killer cells antigen CD94 |
| OID20596 | P08727 | KRT19 | Keratin, type I cytoskeletal 19 |
| OID20669 | Q16719 | KYNU | Kynureninase |
| OID20737 | Q6GTX8 | LAIR1 | Leukocyte-associated immunoglobulin-like receptor 1 |
| OID20769 | Q16363 | LAMA4 | Laminin subunit alpha-4 |
| OID20638 | Q9UQV4 | LAMP3 | Lysosome-associated membrane glycoprotein 3 |
| OID20436 | P28838 | LAP3 | Cytosol aminopeptidase |
| OID20640 | O43561 | LAT | Linker for activation of T-cells family member 1 |
| OID20748 | P56470 | LGALS4 | Galectin-4 |
| OID20781 | O00182 | LGALS9 | Galectin-9 |
| OID20773 | Q99538 | LGMN | Legumain |
| OID20749 | Q9H008 | LHPP | Phospholysine phosphohistidine inorganic pyrophosphate phosphatase |
| OID20606 | P42702 | LIFR | Leukemia inhibitory factor receptor |
| OID20429 | Q8NHJ6 | LILRB4 | Leukocyte immunoglobulin-like receptor subfamily B member 4 |
| OID20438 | Q6UXK5 | LRRN1 | Leucine-rich repeat neuronal protein 1 |
| OID20530 | P33241 | LSP1 | Lymphocyte-specific protein 1 |
| OID20586 | P01374 | LTA | Lymphotoxin-alpha |
| OID20768 | P36941 | LTBR | Tumor necrosis factor receptor superfamily member 3 |
| OID20434 | Q8WV07 | LTO1 | Protein LTO1 homolog |
| OID20708 | Q14210 | LY6D | Lymphocyte antigen 6D |
| OID20580 | O60449 | LY75 | Lymphocyte antigen 75 |
| OID20670 | Q9HBG7 | LY9 | T-lymphocyte surface antigen Ly-9 |
| OID20707 | P55145 | MANF | Mesencephalic astrocyte-derived neurotrophic factor |
| OID20552 | P52564 | MAP2K6 | Dual specificity mitogen-activated protein kinase kinase 6 |
| OID20557 | P45984 | MAPK9 | Mitogen-activated protein kinase 9 |
| OID20767 | O00339 | MATN2 | Matrilin-2 |
| OID20746 | Q96KG7 | MEGF10 | Multiple epidermal growth factor-like domains protein 10 |
| OID20753 | Q9NQ76 | MEPE | Matrix extracellular phosphoglycoprotein |
| OID20657 | Q12866 | MERTK | Tyrosine-protein kinase Mer |
| OID20519 | Q6UB28 | METAP1D | Methionine aminopeptidase 1D, mitochondrial |
| OID20711 | Q99685 | MGLL | Monoglyceride lipase |
| OID20588 | P16455 | MGMT | Methylated-DNA–protein-cysteine methyltransferase |
| OID20593 | Q29980_Q29983 | MICB_MICA | MHC class I polypeptide-related sequence A_MHC class I polypeptide-related s |
| OID20482 | Q7Z6M3 | MILR1 | Allergin-1 |
| OID20541 | P12872 | MLN | Promotilin |
| OID20672 | P03956 | MMP1 | Interstitial collagenase |
| OID20687 | P09238 | MMP10 | Stromelysin-2 |
| OID20779 | O95866 | MPIG6B | Megakaryocyte and platelet inhibitory receptor G6b |
| OID20459 | Q03426 | MVK | Mevalonate kinase |
| OID20451 | Q13459 | MYO9B | Unconventional myosin-IXb |
| OID20732 | Q8WU39 | MZB1 | Marginal zone B- and B1-cell-specific protein |
| OID20442 | O60934 | NBN | Nibrin |
| OID20567 | P19878 | NCF2 | Neutrophil cytosol factor 2 |
| OID20683 | O43639 | NCK2 | Cytoplasmic protein NCK2 |
| OID20448 | Q969V3 | NCLN | Nicalin |
| OID20566 | O76036 | NCR1 | Natural cytotoxicity triggering receptor 1 |
| OID20706 | Q99435 | NELL2 | Protein kinase C-binding protein NELL2 |
| OID20634 | O94856 | NFASC | Neurofascin |
| OID20545 | O95644 | NFATC1 | Nuclear factor of activated T-cells, cytoplasmic 1 |
| OID20470 | Q12968 | NFATC3 | Nuclear factor of activated T-cells, cytoplasmic 3 |
| OID20789 | Q13232 | NME3 | Nucleoside diphosphate kinase 3 |
| OID20568 | P23582 | NPPC | C-type natriuretic peptide |
| OID20465 | Q99748 | NRTN | Neurturin |
| OID20553 | Q9H0P0 | NT5C3A | Cytosolic 5′-nucleotidase 3A |
| OID20591 | P20783 | NTF3 | Neurotrophin-3 |
| OID20510 | Q9Y5A7 | NUB1 | NEDD8 ultimate buster 1 |
| OID20623 | Q9Y266 | NUDC | Nuclear migration protein nudC |
| OID20725 | Q99983 | OMD | Osteomodulin |
| OID20776 | Q8IYS5 | OSCAR | Osteoclast-associated immunoglobulin-like receptor |
| OID20574 | P13725 | OSM | Oncostatin-M |
| OID20467 | Q9Y2J8 | PADI2 | Protein-arginine deiminase type-2 |
| OID20421 | Q13219 | PAPPA | Pappalysin-1 |
| OID20500 | P09874 | PARP1 | Poly [ADP-ribose] polymerase 1 |
| OID20614 | Q08174 | PCDH1 | protocadherin-1 |
| OID20741 | P01127 | PDGFB | Platelet-derived growth factor subunit B |
| OID20729 | Q9NR12 | PDLIM7 | PDZ and LIM domain protein 7 |
| OID20673 | P49763 | PGF | Placenta growth factor |
| OID20587 | Q6ZUJ8 | PIK3AP1 | Phosphoinositide 3-kinase adapter protein 1 |
| OID20682 | P30613 | PKLR | Pyruvate kinase PKLR |
| OID20685 | P47712 | PLA2G4A | Cytosolic phospholipase A2 |
| OID20764 | Q03405 | PLAUR | Urokinase plasminogen activator surface receptor |
| OID20597 | Q9HCM2 | PLXNA4 | Plexin-A4 |
| OID20775 | P54317 | PNLIPRP2 | Pancreatic lipase-related protein 2 |
| OID20445 | Q8TCS8 | PNPT1 | Polyribonucleotide nucleotidyltransferase 1, mitochondrial |
| OID20777 | Q15166 | PON3 | Serum paraoxonase/lactonase 3 |
| OID20643 | Q96SB3 | PPP1R9B | Neurabin-2 |
| OID20483 | P30048 | PRDX3 | Thioredoxin-dependent peroxide reductase, mitochondrial |
| OID20644 | P30044 | PRDX5 | Peroxiredoxin-5, mitochondrial |
| OID20439 | Q9HCU5 | PREB | Prolactin regulatory element-binding protein |
| OID20766 | P51888 | PRELP | Prolargin |
| OID20444 | Q9Y478 | PRKAB1 | 5′-AMP-activated protein kinase subunit beta-1 |
| OID20468 | Q04759 | PRKCQ | Protein kinase C theta type |
| OID20543 | P58294 | PROK1 | Prokineticin-1 |
| OID20763 | Q16651 | PRSS8 | Prostasin |
| OID20534 | O75475 | PSIP1 | PC4 and SFRS1-interacting protein |
| OID20549 | Q9BT73 | PSMG3 | Proteasome assembly chaperone 3 |
| OID20511 | O60542 | PSPN | Persephin |
| OID20571 | Q03431 | PTH1R | Parathyroid hormone/parathyroid hormone-related peptide receptor |
| OID20717 | P29350 | PTPN6 | Tyrosine-protein phosphatase non-receptor type 6 |
| OID20598 | P28827 | PTPRM | protein tyrosine phosphatase receptor type M |
| OID20570 | P26022 | PTX3 | Pentraxin-related protein PTX3 |
| OID20425 | Q96AX2 | RAB37 | Ras-related protein Rab-37 |
| OID20608 | P20340 | RAB6A | Ras-related protein Rab-6A |
| OID20447 | Q5R372 | RABGAP1L | Rab GTPase-activating protein 1-like |
| OID20784 | Q9BYZ8 | REG4 | Regenerating islet-derived protein 4 |
| OID20454 | P57771 | RGS8 | Regulator of G-protein signaling 8 |
| OID20734 | Q9Y6N7 | ROBO1 | Roundabout homolog 1 |
| OID20536 | Q8IVG5 | SAMD9L | Sterile alpha motif domain-containing protein 9-like |
| OID20728 | Q8WXD2 | SCG3 | Secretogranin-3 |
| OID20688 | P11684 | SCGB1A1 | Uteroglobin |
| OID20723 | Q96PL1 | SCGB3A2 | Secretoglobin family 3A member 2 |
| OID20457 | O76038 | SCGN | Secretagogin |
| OID20542 | Q12765 | SCRN1 | Secernin-1 |
| OID20560 | Q14242 | SELPLG | Selectin P Ligand |
| OID20630 | P50452 | SERPINB8 | Serpin B8 |
| OID20514 | O60880 | SH2D1A | SH2 domain-containing protein 1A |
| OID20714 | P34896 | SHMT1 | Serine hydroxymethyltransferase, cytosolic |
| OID20690 | Q9BZZ2 | SIGLEC1 | Sialoadhesin |
| OID20537 | Q96LC7 | SIGLEC10 | Sialic acid-binding Ig-like lectin 10 |
| OID20739 | O00241 | SIRPB1 | Signal-regulatory protein beta-1 |
| OID20462 | Q9Y3P8 | SIT1 | Signaling threshold-regulating transmembrane adapter 1 |
| OID20761 | O75563 | SKAP2 | Src kinase-associated phosphoprotein 2 |
| OID20496 | Q13291 | SLAMF1 | Signaling lymphocytic activation molecule |
| OID20602 | Q9NQ25 | SLAMF7 | SLAM family member 7 |
| OID20539 | Q6ZMH5 | SLC39A5 | Zinc transporter ZIP5 |
| OID20730 | Q9H3U7 | SMOC2 | SPARC-related modular calcium-binding protein 2 |
| OID20676 | Q92484 | SMPDL3A | Acid sphingomyelinase-like phosphodiesterase 3a |
| OID20509 | O60575 | SPINK4 | Serine protease inhibitor Kazal-type 4 |
| OID20720 | O43291 | SPINT2 | Kunitz-type protease inhibitor 2 |
| OID20759 | Q9HCB6 | SPON1 | Spondin-1 |
| OID20475 | O43597 | SPRY2 | Protein sprouty homolog 2 |
| OID20502 | P78362 | SRPK2 | SRSF protein kinase 2 |
| OID20531 | Q9UNK0 | STX8 | Syntaxin-8 |
| OID20508 | Q06520 | SULT2A1 | Sulfotransferase 2A1 |
| OID20474 | Q92844 | TANK | TRAF family member-associated NF-kappa-B activator |
| OID20521 | Q92609 | TBC1D5 | TBC1 domain family member 5 |
| OID20787 | Q03403 | TFF2 | Trefoil factor 2 |
| OID20600 | P01135 | TGFA | Protransforming growth factor alpha |
| OID20621 | P01137 | TGFB1 | Transforming growth factor beta-1 proprotein |
| OID20684 | P35625 | TIMP3 | Metalloproteinase inhibitor 3 |
| OID20612 | O15455 | TLR3 | Toll-like receptor 3 |
| OID20473 | P01375 | TNF | Tumor necrosis factor |
| OID20433 | O95379 | TNFAIP8 | Tumor necrosis factor alpha-induced protein 8 |
| OID20646 | Q9Y6Q6 | TNFRSF11A | Tumor necrosis factor receptor superfamily member 11A |
| OID20735 | O00300 | TNFRSF11B | Tumor necrosis factor receptor superfamily member 11B |
| OID20702 | O14836 | TNFRSF13B | Tumor necrosis factor receptor superfamily member 13B |
| OID20480 | Q96RJ3 | TNFRSF13C | Tumor necrosis factor receptor superfamily member 13C |
| OID20783 | Q92956 | TNFRSF14 | Tumor necrosis factor receptor superfamily member 14 |
| OID20653 | P43489 | TNFRSF4 | Tumor necrosis factor receptor superfamily member 4 |
| OID20611 | P50591 | TNFSF10 | Tumor necrosis factor ligand superfamily member 10 |
| OID20592 | O14788 | TNFSF11 | Tumor necrosis factor ligand superfamily member 11 |
| OID20624 | O43508 | TNFSF12 | Tumor necrosis factor ligand superfamily member 12 |
| OID20733 | O75888 | TNFSF13 | Tumor necrosis factor ligand superfamily member 13 |
| OID20750 | O14773 | TPP1 | Tripeptidyl-peptidase 1 |
| OID20642 | Q15661 | TPSAB1 | Tryptase alpha/beta-1 |
| OID20476 | P13693 | TPT1 | Translationally-controlled tumor protein |
| OID20507 | Q12933 | TRAF2 | TNF receptor-associated factor 2 |
| OID20731 | Q9NZC2 | TREM2 | Triggering receptor expressed on myeloid cells 2 |
| OID20533 | P19474 | TRIM21 | E3 ubiquitin-protein ligase TRIM21 |
| OID20526 | Q9C035 | TRIM5 | Tripartite motif-containing protein 5 |
| OID20516 | Q7L8A9 | VASH1 | Tubulinyl-Tyr carboxypeptidase 1 |
| OID20650 | P15692 | VEGFA | Vascular endothelial growth factor A |
| OID20662 | O43915 | VEGFD | Vascular endothelial growth factor D |
| OID20479 | P42768 | WAS | Actin nucleation-promoting factor WAS |
| OID20785 | Q8TEU8 | WFIKKN2 | WAP, Kazal, immunoglobulin, Kunitz and NTR domain-containing protein 2 |
| OID20460 | O14904 | WNT9A | Protein Wnt-9a |
| OID20478 | Q7Z739 | YTHDF3 | YTH domain-containing family protein 3 |
