## Supplemental Table 2 for "The maternal inflammatory proteome during pregnancy and its role in predicting the risk of spontaneous preterm birth"

**Supplementary Table 2. Protein Missingness and QC warning**

| **OlinkID** | **UniProt** | **Assay** | **MissingFreq (%)** | **QC_Warning** |
| --- | --- | --- | --- | --- |
| OID20435 | O43707 | ACTN4 | 64.77 | PASS |
| OID20645 | P00813 | ADA | 0 | PASS |
| OID20651 | O75077 | ADAM23 | 0 | PASS |
| OID20755 | Q9UHX3 | ADGRE2 | 0 | PASS |
| OID20756 | Q15109 | AGER | 0 | PASS |
| OID20786 | O00468 | AGRN | 0 | PASS |
| OID20658 | O00253 | AGRP | 0 | PASS |
| OID20512 | P30838 | ALDH3A1 | 21.59 | PASS |
| OID20437 | Q9NP70 | AMBN | 0 | PASS |
| OID20492 | Q9BXJ7 | AMN | 27.27 | PASS |
| OID20740 | Q15389 | ANGPT1 | 0 | PASS |
| OID20726 | Q9UKU9 | ANGPTL2 | 0 | PASS |
| OID20703 | Q9BY76 | ANGPTL4 | 0 | PASS |
| OID20583 | P50995 | ANXA11 | 0 | PASS |
| OID20452 | P19801 | AOC1 | 0 | PASS |
| OID20554 | Q9NZN5 | ARHGEF12 | 2.27 | PASS |
| OID20432 | P27540 | ARNT | 68.75 | PASS |
| OID20446 | Q5T4W7 | ARTN | 55.11 | PASS |
| OID20760 | Q9UII2 | ATP5IF1 | 0 | PASS |
| OID20582 | O15169 | AXIN1 | 0 | PASS |
| OID20780 | P15291 | B4GALT1 | 0 | PASS |
| OID20505 | O14867 | BACH1 | 12.5 | PASS |
| OID20594 | Q8NDB2 | BANK1 | 0 | PASS |
| OID20513 | O43521-2 | BCL2L11 | NA | EXCLUDED |
| OID20558 | P11274 | BCR | 0 | PASS |
| OID20517 | P55957 | BID | NA | EXCLUDED |
| OID20626 | P35613 | BSG | 0 | PASS |
| OID20713 | Q7KYR7 | BTN2A1 | 0 | PASS |
| OID20540 | P78410 | BTN3A2 | 0 | PASS |
| OID20654 | P02745 | C1QA | 0 | PASS |
| OID20555 | P42575 | CASP2 | 3.41 | PASS |
| OID20668 | P51671 | CCL11 | 0 | PASS |
| OID20655 | Q99616 | CCL13 | 0 | PASS |
| OID20745 | Q92583 | CCL17 | 0 | PASS |
| OID20671 | P78556 | CCL20 | 0 | PASS |
| OID20686 | O00585 | CCL21 | 0 | PASS |
| OID20765 | O00626 | CCL22 | 0 | PASS |
| OID20693 | P55773 | CCL23 | 0 | PASS |
| OID20772 | O00175 | CCL24 | 0 | PASS |
| OID20674 | O15444 | CCL25 | 0 | PASS |
| OID20546 | Q9Y258 | CCL26 | 0 | PASS |
| OID20569 | Q9NRJ3 | CCL28 | 0 | PASS |
| OID20610 | P10147 | CCL3 | 0 | PASS |
| OID20695 | P13236 | CCL4 | 0 | PASS |
| OID20523 | P80098 | CCL7 | 0 | PASS |
| OID20709 | P29279 | CCN2 | 0 | PASS |
| OID20647 | O95971 | CD160 | 0 | PASS |
| OID20603 | P41217 | CD200 | 0 | PASS |
| OID20595 | Q8TD46 | CD200R1 | 0 | PASS |
| OID20637 | P20273 | CD22 | 0 | PASS |
| OID20628 | Q9BZW8 | CD244 | 0 | PASS |
| OID20680 | Q5ZPR3 | CD276 | 0 | PASS |
| OID20584 | P01730 | CD4 | 0 | PASS |
| OID20724 | P25942 | CD40 | 0 | PASS |
| OID20616 | P29965 | CD40LG | 0 | PASS |
| OID20692 | P09326 | CD48 | 0 | PASS |
| OID20716 | P19256 | CD58 | 0 | PASS |
| OID20649 | P30203 | CD6 | 0 | PASS |
| OID20599 | P32970 | CD70 | 0 | PASS |
| OID20635 | P40259 | CD79B | 0 | PASS |
| OID20565 | Q01151 | CD83 | 0 | PASS |
| OID20607 | Q9UIB8 | CD84 | 0 | PASS |
| OID20754 | Q4KMG0 | CDON | 0 | PASS |
| OID20667 | Q15517 | CDSN | 0 | PASS |
| OID20548 | Q3KPI0 | CEACAM21 | 0 | PASS |
| OID20441 | Q9UPV0 | CEP164 | 89.77 | PASS |
| OID20771 | Q9BU40 | CHRDL1 | 0 | PASS |
| OID20617 | Q07065 | CKAP4 | 0 | PASS |
| OID20721 | P12532 | CKMT1A_CKMT1B | 0 | PASS |
| OID20573 | Q9UMR7 | CLEC4A | 0 | PASS |
| OID20547 | Q8WTT0 | CLEC4C | 0 | PASS |
| OID20609 | Q8WXI8 | CLEC4D | 0 | PASS |
| OID20590 | Q6UXB4 | CLEC4G | 0 | PASS |
| OID20636 | Q9BXN2 | CLEC7A | 1.14 | PASS |
| OID20559 | Q9UDT6 | CLIP2 | 0 | PASS |
| OID20664 | Q9H4D0 | CLSTN2 | 0 | PASS |
| OID20556 | Q9UHC6 | CNTNAP2 | 0 | PASS |
| OID20550 | P20849 | COL9A1 | 0 | PASS |
| OID20738 | Q5KU26 | COLEC12 | 0 | PASS |
| OID20751 | Q6UXH1 | CRELD2 | 0 | PASS |
| OID20747 | P24387 | CRHBP | 0 | PASS |
| OID20701 | Q9NZV1 | CRIM1 | 0 | PASS |
| OID20742 | P46109 | CRKL | 0 | PASS |
| OID20699 | O75462 | CRLF1 | 0 | PASS |
| OID20719 | P09603 | CSF1 | 0 | PASS |
| OID20491 | P09919 | CSF3 | 0 | PASS |
| OID20704 | O76096 | CST7 | 0 | PASS |
| OID20752 | Q99895 | CTRC | 0 | PASS |
| OID20679 | P53634 | CTSC | 0 | PASS |
| OID20660 | P43234 | CTSO | 0 | PASS |
| OID20561 | P78310 | CXADR | 0 | PASS |
| OID20762 | P09341 | CXCL1 | 0 | PASS |
| OID20697 | P02778 | CXCL10 | 0 | PASS |
| OID20464 | P48061 | CXCL12 | 0 | PASS |
| OID20458 | O95715 | CXCL14 | 94.32 | PASS |
| OID20622 | Q6UXB2 | CXCL17 | 0 | PASS |
| OID20788 | P19876 | CXCL3 | 0 | PASS |
| OID20613 | P80162 | CXCL6 | 0 | PASS |
| OID20631 | P10145 | CXCL8 | 0 | PASS |
| OID20675 | Q07325 | CXCL9 | 0 | PASS |
| OID20774 | Q14118 | DAG1 | 0 | PASS |
| OID20524 | Q9UN19 | DAPP1 | 0 | PASS |
| OID20681 | Q9UJU6 | DBNL | 0 | PASS |
| OID20579 | Q16698 | DECR1 | 0 | PASS |
| OID20620 | O00273 | DFFA | 0 | PASS |
| OID20466 | Q13574 | DGKZ | 82.95 | PASS |
| OID20627 | O60884 | DNAJA2 | 0 | PASS |
| OID20712 | Q8NFT8 | DNER | 0 | PASS |
| OID20744 | O43598 | DNPH1 | 0 | PASS |
| OID20527 | Q8N608 | DPP10 | 0 | PASS |
| OID20525 | Q9UNE0 | EDAR | 0 | PASS |
| OID20698 | P01133 | EGF | 0 | PASS |
| OID20572 | Q9GZT9 | EGLN1 | 0 | PASS |
| OID20625 | Q04637 | EIF4G1 | 0 | PASS |
| OID20423 | P63241 | EIF5A | 58.52 | PASS |
| OID20497 | Q8N8S7 | ENAH | 0 | PASS |
| OID20736 | Q9UJA9 | ENPP5 | 0 | PASS |
| OID20689 | Q6UWV6 | ENPP7 | 0 | PASS |
| OID20743 | P16422 | EPCAM | 0 | PASS |
| OID20677 | P21709 | EPHA1 | 0 | PASS |
| OID20522 | P01588 | EPO | 0 | PASS |
| OID20705 | P21860 | ERBB3 | 0 | PASS |
| OID20758 | Q9NQ30 | ESM1 | 0 | PASS |
| OID20691 | P25116 | F2R | 0 | PASS |
| OID20778 | P07148 | FABP1 | 0 | PASS |
| OID20501 | Q0Z7S8 | FABP9 | 0 | PASS |
| OID20665 | P48023 | FASLG | 0 | PASS |
| OID20564 | P24071 | FCAR | 0 | PASS |
| OID20639 | Q96LA5 | FCRL2 | 0 | PASS |
| OID20443 | Q96P31 | FCRL3 | 2.84 | PASS |
| OID20529 | Q6DN72 | FCRL6 | 0 | PASS |
| OID20715 | O95750 | FGF19 | 0 | PASS |
| OID20503 | P09038 | FGF2 | 32.95 | PASS |
| OID20490 | P12034 | FGF5 | 0 | PASS |
| OID20722 | Q9Y3D6 | FIS1 | 0 | PASS |
| OID20618 | P68106 | FKBP1B | 2.27 | PASS |
| OID20661 | P49771 | FLT3LG | 0 | PASS |
| OID20515 | Q12778 | FOXO1 | 0 | PASS |
| OID20770 | P19883 | FST | 0 | PASS |
| OID20782 | O95633 | FSTL3 | 0 | PASS |
| OID20463 | Q96DB9 | FXYD5 | 0.57 | PASS |
| OID20727 | P22466 | GAL | 0 | PASS |
| OID20471 | Q14435 | GALNT3 | 0 | PASS |
| OID20489 | P32456 | GBP2 | 55.68 | PASS |
| OID20710 | Q9HC38 | GLOD4 | 0 | PASS |
| OID20641 | P36959 | GMPR | 0 | PASS |
| OID20551 | Q9HD26 | GOPC | 0 | PASS |
| OID20663 | P12544 | GZMA | 0 | PASS |
| OID20604 | P10144 | GZMB | 94.32 | PASS |
| OID20648 | P14317 | HCLS1 | 0 | PASS |
| OID20589 | O94992 | HEXIM1 | 0 | PASS |
| OID20656 | P14210 | HGF | 0 | PASS |
| OID20520 | P01903 | HLA-DRA | 0 | PASS |
| OID20532 | P13747 | HLA-E | 92.61 | PASS |
| OID20615 | P37235 | HPCAL1 | 0 | PASS |
| OID20575 | P28845 | HSD11B1 | 0 | PASS |
| OID20718 | P0DMV8 | HSPA1A | 0 | PASS |
| OID20484 | Q05084 | ICA1 | 0.57 | PASS |
| OID20578 | Q14773 | ICAM4 | 0 | PASS |
| OID20619 | P22304 | IDS | 0 | PASS |
| OID20495 | P01579 | IFNG | 1.7 | PASS |
| OID20678 | P15260 | IFNGR1 | 0 | PASS |
| OID20506 | Q8IU57 | IFNLR1 | 0 | PASS |
| OID20544 | Q9Y6K9 | IKBKG | 0 | PASS |
| OID20431 | P22301 | IL10 | 0 | PASS |
| OID20420 | Q13651 | IL10RA | 27.84 | PASS |
| OID20696 | Q08334 | IL10RB | 0 | PASS |
| OID20455 | P20809 | IL11 | 40.91 | PASS |
| OID20666 | P29460 | IL12B | 0 | PASS |
| OID20486 | P42701 | IL12RB1 | 1.14 | PASS |
| OID20430 | P35225 | IL13 | 32.39 | PASS |
| OID20562 | P40933 | IL15 | 0 | PASS |
| OID20498 | Q13261 | IL15RA | 92.61 | PASS |
| OID20633 | Q14005 | IL16 | 0 | PASS |
| OID20469 | Q16552 | IL17A | 0 | PASS |
| OID20477 | Q9P0M4 | IL17C | 0 | PASS |
| OID20481 | Q8TAD2 | IL17D | 0 | PASS |
| OID20456 | Q96PD4 | IL17F | 27.27 | PASS |
| OID20585 | Q9NRM6 | IL17RB | 0 | PASS |
| OID20694 | Q14116 | IL18 | 0 | PASS |
| OID20652 | Q13478 | IL18R1 | 0 | PASS |
| OID20488 | P01583 | IL1A | 82.39 | PASS |
| OID20427 | P01584 | IL1B | 0 | PASS |
| OID20757 | P27930 | IL1R2 | 0 | PASS |
| OID20487 | Q9HB29 | IL1RL2 | 0 | PASS |
| OID20700 | P18510 | IL1RN | 0 | PASS |
| OID20419 | P60568 | IL2 | 4.55 | PASS |
| OID20453 | Q9NYY1 | IL20 | 77.27 | PASS |
| OID20422 | Q9UHF4 | IL20RA | 16.48 | PASS |
| OID20449 | Q8N6P7 | IL22RA1 | 3.41 | PASS |
| OID20440 | Q13007 | IL24 | 91.48 | PASS |
| OID20450 | P14784 | IL2RB | 34.66 | PASS |
| OID20605 | P24001 | IL32 | 0 | PASS |
| OID20428 | O95760 | IL33 | 82.39 | PASS |
| OID20461 | P26951 | IL3RA | 6.25 | PASS |
| OID20426 | P05112 | IL4 | 97.73 | PASS |
| OID20576 | P24394 | IL4R | 0 | PASS |
| OID20472 | P05113 | IL5 | 77.84 | PASS |
| OID20601 | Q01344 | IL5RA | 0 | PASS |
| OID20563 | P05231 | IL6 | 0 | PASS |
| OID20535 | P13232 | IL7 | 0 | PASS |
| OID20485 | P51617 | IRAK1 | 0 | PASS |
| OID20577 | Q9NWZ3 | IRAK4 | 0 | PASS |
| OID20538 | B1AKI9 | ISM1 | 0 | PASS |
| OID20581 | Q9UKX5 | ITGA11 | 0 | PASS |
| OID20528 | P23229 | ITGA6 | 11.36 | PASS |
| OID20493 | P18564 | ITGB6 | 0 | PASS |
| OID20504 | O43736 | ITM2A | 0 | PASS |
| OID20494 | P01591 | JCHAIN | 0 | PASS |
| OID20424 | P05412 | JUN | 6.25 | PASS |
| OID20629 | Q12918 | KLRB1 | 0 | PASS |
| OID20632 | Q13241 | KLRD1 | 0 | PASS |
| OID20596 | P08727 | KRT19 | 0 | PASS |
| OID20669 | Q16719 | KYNU | 0 | PASS |
| OID20737 | Q6GTX8 | LAIR1 | 0 | PASS |
| OID20769 | Q16363 | LAMA4 | 0 | PASS |
| OID20638 | Q9UQV4 | LAMP3 | 0 | PASS |
| OID20436 | P28838 | LAP3 | 1.14 | PASS |
| OID20640 | O43561 | LAT | 0 | PASS |
| OID20748 | P56470 | LGALS4 | 0 | PASS |
| OID20781 | O00182 | LGALS9 | 0 | PASS |
| OID20773 | Q99538 | LGMN | 0 | PASS |
| OID20749 | Q9H008 | LHPP | 0 | PASS |
| OID20606 | P42702 | LIFR | 0 | PASS |
| OID20429 | Q8NHJ6 | LILRB4 | 0 | PASS |
| OID20438 | Q6UXK5 | LRRN1 | 0 | PASS |
| OID20530 | P33241 | LSP1 | 0 | PASS |
| OID20586 | P01374 | LTA | 0.57 | PASS |
| OID20768 | P36941 | LTBR | 0 | PASS |
| OID20434 | Q8WV07 | LTO1 | 46.59 | PASS |
| OID20708 | Q14210 | LY6D | 0 | PASS |
| OID20580 | O60449 | LY75 | 0 | PASS |
| OID20670 | Q9HBG7 | LY9 | 0 | PASS |
| OID20707 | P55145 | MANF | 0 | PASS |
| OID20552 | P52564 | MAP2K6 | 0 | PASS |
| OID20557 | P45984 | MAPK9 | 0 | PASS |
| OID20767 | O00339 | MATN2 | 0 | PASS |
| OID20746 | Q96KG7 | MEGF10 | 0 | PASS |
| OID20753 | Q9NQ76 | MEPE | 0 | PASS |
| OID20657 | Q12866 | MERTK | 0 | PASS |
| OID20519 | Q6UB28 | METAP1D | 0 | PASS |
| OID20711 | Q99685 | MGLL | NA | EXCLUDED |
| OID20588 | P16455 | MGMT | 0 | PASS |
| OID20593 | Q29980_Q29983 | MICB_MICA | 4.55 | PASS |
| OID20482 | Q7Z6M3 | MILR1 | 0 | PASS |
| OID20541 | P12872 | MLN | 0 | PASS |
| OID20672 | P03956 | MMP1 | 0 | PASS |
| OID20687 | P09238 | MMP10 | 0 | PASS |
| OID20779 | O95866 | MPIG6B | 0 | PASS |
| OID20459 | Q03426 | MVK | 0 | PASS |
| OID20451 | Q13459 | MYO9B | 3.98 | PASS |
| OID20732 | Q8WU39 | MZB1 | 0 | PASS |
| OID20442 | O60934 | NBN | 0 | PASS |
| OID20567 | P19878 | NCF2 | 0 | PASS |
| OID20683 | O43639 | NCK2 | 0 | PASS |
| OID20448 | Q969V3 | NCLN | 71.59 | PASS |
| OID20566 | O76036 | NCR1 | 0 | PASS |
| OID20706 | Q99435 | NELL2 | 0 | PASS |
| OID20634 | O94856 | NFASC | 5.68 | PASS |
| OID20545 | O95644 | NFATC1 | 0 | PASS |
| OID20470 | Q12968 | NFATC3 | 85.8 | PASS |
| OID20789 | Q13232 | NME3 | 0 | PASS |
| OID20568 | P23582 | NPPC | 0 | PASS |
| OID20465 | Q99748 | NRTN | 34.09 | PASS |
| OID20553 | Q9H0P0 | NT5C3A | 0 | PASS |
| OID20591 | P20783 | NTF3 | 0 | PASS |
| OID20510 | Q9Y5A7 | NUB1 | 0 | PASS |
| OID20623 | Q9Y266 | NUDC | 0 | PASS |
| OID20725 | Q99983 | OMD | 0 | PASS |
| OID20776 | Q8IYS5 | OSCAR | 0 | PASS |
| OID20574 | P13725 | OSM | 0 | PASS |
| OID20467 | Q9Y2J8 | PADI2 | 19.89 | PASS |
| OID20421 | Q13219 | PAPPA | 0 | PASS |
| OID20500 | P09874 | PARP1 | 83.52 | PASS |
| OID20614 | Q08174 | PCDH1 | 0 | PASS |
| OID20741 | P01127 | PDGFB | 0 | PASS |
| OID20729 | Q9NR12 | PDLIM7 | 0 | PASS |
| OID20673 | P49763 | PGF | 0 | PASS |
| OID20587 | Q6ZUJ8 | PIK3AP1 | 0 | PASS |
| OID20682 | P30613 | PKLR | 0 | PASS |
| OID20685 | P47712 | PLA2G4A | 0 | PASS |
| OID20764 | Q03405 | PLAUR | 0 | PASS |
| OID20597 | Q9HCM2 | PLXNA4 | 0 | PASS |
| OID20775 | P54317 | PNLIPRP2 | 0 | PASS |
| OID20445 | Q8TCS8 | PNPT1 | 36.93 | PASS |
| OID20777 | Q15166 | PON3 | 0 | PASS |
| OID20643 | Q96SB3 | PPP1R9B | 0 | PASS |
| OID20483 | P30048 | PRDX3 | 13.07 | PASS |
| OID20644 | P30044 | PRDX5 | 0 | PASS |
| OID20439 | Q9HCU5 | PREB | 75 | PASS |
| OID20766 | P51888 | PRELP | 0 | PASS |
| OID20444 | Q9Y478 | PRKAB1 | 1.14 | PASS |
| OID20468 | Q04759 | PRKCQ | 59.09 | PASS |
| OID20543 | P58294 | PROK1 | 0 | PASS |
| OID20763 | Q16651 | PRSS8 | 0 | PASS |
| OID20534 | O75475 | PSIP1 | 0 | PASS |
| OID20549 | Q9BT73 | PSMG3 | 0 | PASS |
| OID20511 | O60542 | PSPN | 12.5 | PASS |
| OID20571 | Q03431 | PTH1R | 0 | PASS |
| OID20717 | P29350 | PTPN6 | 0 | PASS |
| OID20598 | P28827 | PTPRM | 0 | PASS |
| OID20570 | P26022 | PTX3 | 0 | PASS |
| OID20425 | Q96AX2 | RAB37 | 1.14 | PASS |
| OID20608 | P20340 | RAB6A | 11.93 | PASS |
| OID20447 | Q5R372 | RABGAP1L | 0 | PASS |
| OID20784 | Q9BYZ8 | REG4 | 0 | PASS |
| OID20454 | P57771 | RGS8 | 81.82 | PASS |
| OID20734 | Q9Y6N7 | ROBO1 | 0 | PASS |
| OID20536 | Q8IVG5 | SAMD9L | 0 | PASS |
| OID20728 | Q8WXD2 | SCG3 | 0 | PASS |
| OID20688 | P11684 | SCGB1A1 | 0 | PASS |
| OID20723 | Q96PL1 | SCGB3A2 | 0 | PASS |
| OID20457 | O76038 | SCGN | 1.7 | PASS |
| OID20542 | Q12765 | SCRN1 | 0.57 | PASS |
| OID20560 | Q14242 | SELPLG | 0 | PASS |
| OID20630 | P50452 | SERPINB8 | 0 | PASS |
| OID20514 | O60880 | SH2D1A | 0 | PASS |
| OID20714 | P34896 | SHMT1 | 0 | PASS |
| OID20690 | Q9BZZ2 | SIGLEC1 | 0 | PASS |
| OID20537 | Q96LC7 | SIGLEC10 | 0 | PASS |
| OID20739 | O00241 | SIRPB1 | 0 | PASS |
| OID20462 | Q9Y3P8 | SIT1 | 0.57 | PASS |
| OID20761 | O75563 | SKAP2 | 0 | PASS |
| OID20496 | Q13291 | SLAMF1 | 26.14 | PASS |
| OID20602 | Q9NQ25 | SLAMF7 | 0 | PASS |
| OID20539 | Q6ZMH5 | SLC39A5 | 0 | PASS |
| OID20730 | Q9H3U7 | SMOC2 | 0 | PASS |
| OID20676 | Q92484 | SMPDL3A | 0 | PASS |
| OID20509 | O60575 | SPINK4 | 9.66 | PASS |
| OID20720 | O43291 | SPINT2 | 0 | PASS |
| OID20759 | Q9HCB6 | SPON1 | 0 | PASS |
| OID20475 | O43597 | SPRY2 | 0 | PASS |
| OID20502 | P78362 | SRPK2 | 7.95 | PASS |
| OID20531 | Q9UNK0 | STX8 | 5.68 | PASS |
| OID20508 | Q06520 | SULT2A1 | 2.84 | PASS |
| OID20474 | Q92844 | TANK | 3.98 | PASS |
| OID20521 | Q92609 | TBC1D5 | 3.41 | PASS |
| OID20787 | Q03403 | TFF2 | 0 | PASS |
| OID20600 | P01135 | TGFA | 0 | PASS |
| OID20621 | P01137 | TGFB1 | 0 | PASS |
| OID20684 | P35625 | TIMP3 | 0 | PASS |
| OID20612 | O15455 | TLR3 | 0 | PASS |
| OID20473 | P01375 | TNF | 0 | PASS |
| OID20433 | O95379 | TNFAIP8 | 1.14 | PASS |
| OID20646 | Q9Y6Q6 | TNFRSF11A | 0 | PASS |
| OID20735 | O00300 | TNFRSF11B | 0 | PASS |
| OID20702 | O14836 | TNFRSF13B | 0 | PASS |
| OID20480 | Q96RJ3 | TNFRSF13C | 0 | PASS |
| OID20783 | Q92956 | TNFRSF14 | 0 | PASS |
| OID20653 | P43489 | TNFRSF4 | 0 | PASS |
| OID20611 | P50591 | TNFSF10 | 0 | PASS |
| OID20592 | O14788 | TNFSF11 | 0 | PASS |
| OID20624 | O43508 | TNFSF12 | 0 | PASS |
| OID20733 | O75888 | TNFSF13 | 0 | PASS |
| OID20750 | O14773 | TPP1 | 0 | PASS |
| OID20642 | Q15661 | TPSAB1 | 0 | PASS |
| OID20476 | P13693 | TPT1 | 2.27 | PASS |
| OID20507 | Q12933 | TRAF2 | 0 | PASS |
| OID20731 | Q9NZC2 | TREM2 | 0 | PASS |
| OID20533 | P19474 | TRIM21 | 0 | PASS |
| OID20526 | Q9C035 | TRIM5 | 1.14 | PASS |
| OID20516 | Q7L8A9 | VASH1 | 17.05 | PASS |
| OID20650 | P15692 | VEGFA | 0 | PASS |
| OID20662 | O43915 | VEGFD | 0 | PASS |
| OID20479 | P42768 | WAS | 15.91 | PASS |
| OID20785 | Q8TEU8 | WFIKKN2 | 0 | PASS |
| OID20460 | O14904 | WNT9A | 0 | PASS |
| OID20478 | Q7Z739 | YTHDF3 | 0.57 | PASS |

Proteins shaded orange represent those with missingness greater than 25%.
